# Effects of Maternal Obesity on Fetal Cerebral Glucose Transporter Expression

**DOI:** 10.64898/2026.05.11.723868

**Authors:** Tyler L King, Kevin Prifti, Ruth M Gill, Sarah K England, Antonina I Frolova

**Affiliations:** Department of Pediatrics, Washington University of School of Medicine, St. Louis, MO, USA; Center for Reproductive Health Sciences, Department of Obstetrics and Gynecology, Washington University in St. Louis, St. Louis, MO, USA

**Keywords:** Maternal obesity, fetal brain, neurodevelopment, glucose

## Abstract

Emerging evidence indicates that the maternal in utero environment has enduring effects on offspring neurodevelopment. The obesity epidemic in the United States affects nearly one-third of women before pregnancy, potentially predisposing offspring to harmful developmental conditions. Glucose, the primary energy source for the brain, is highly regulated by facilitative diffusion glucose transporters (GLUTs). However, our understanding of how maternal obesity influences perinatal cerebral glucose metabolism remains limited. We hypothesized that maternal obesity is associated with altered expression of key GLUTs and dysregulated energy-sensing mechanisms in the fetal brain. Female C57BL/6J mice were randomly assigned to either a control diet (CON) or an obesogenic diet (DIO) (60% kcal from fat, 17.5% kcal from sucrose) for 10 weeks, time-mated with control males, and fed their respective diets throughout gestation. At 18.5 days post coitum, fetal brain tissue was collected for protein analysis. DIO diet did not affect litter size, offspring body weight, or brain weight when compared to CON. Whole brain GLUT1 expression was elevated only in female DIO offspring, while GLUT3 and GLUT4 expression was increased in all DIO offspring without modification by sex. However, maternal diet was not associated with differences in the activation of energy regulatory pathways adenosine monophosphate-activated protein kinase (AMPK) or the nutrient-sensing pathway mechanistic target of rapamycin (mTOR) in the fetal brain. These findings suggest that maternal obesogenic diet alters glucose transporter expression in the fetal brain, indicating a potential disruption in cerebral glucose metabolism during critical periods of perinatal development.

## Introduction

In the United States, the prevalence of obesity has increased at a staggering rate and now impacts over one-third of reproductive-age women [1] Maternal obesity is associated with numerous perinatal complications including increased risk for fetal growth abnormalities, congenital anomalies, neonatal injury, and stillbirth [2] Additionally, children born to mothers with obesity have been shown to have increased rates of neurodevelopmental disorders including developmental delay, autism spectrum disorder, attention deficit hyperactivity disorder and cerebral palsy.[3–7] Although the underlying mechanisms contributing to these risks are largely unknown, it has been suggested that the metabolic profile in maternal obesity may be a significant contributing factor in fetal neurodevelopmental programming. There has been extensive preclinical research evaluating how maternal cytokine signaling pathways, microbial dysbiosis and epigenetic regulation affect offspring brain development [8–12]. However, our understanding of the effects maternal obesity has on perinatal cerebral glucose metabolism is surprisingly limited despite glucose being the primary energy source in the brain [13].

Cerebral glucose uptake is tightly regulated to meet the brain’s high energy demands. This process relies on the facilitative diffusion of glucose across the blood-brain barrier and glial cells to reach neuronal tissue [14]. The glucose transport occurs primarily through a family of sodium independent bidirectional facilitative transporters in the solute carrier 2 (*SLC2*) transporter family, also known as glucose transporters (GLUTs) [15]. Several GLUT isoforms have been identified in the brain, with GLUT1 and GLUT3 being the most important in regulating glucose homeostasis [16] GLUT1 is ubiquitously expressed in all tissues and is primarily responsible for basal glucose uptake within the endovascular blood-brain barrier and glial cells in the brain [17–23]. GLUT3 has a high affinity for glucose and is found in tissues with high energy demands. In the brain, it is located on neuronal membranes, mediating the transport of glucose to neurons in energetically-demanding regions such as the cortex, hippocampus, and hypothalamus [24]. GLUT4 is an insulin-sensitive glucose transporter, identified in adipose, cardiac and skeletal muscle tissues [25–27]. In the brain, GLUT4 expression has also been identified on neurons in energy-demanding regions, including the hippocampus, thalamus, cerebellum and cortex [28–31]. Neuronal GLUT4 colocalizes near GLUT3 and insulin receptors [32–34].

Glucose dysregulation in the neonatal period has also been shown to impair cerebral development. For example, episodes of neonatal hypoglycemia are associated with lower executive and visual motor function in preschool-aged children [35]. Despite the known detrimental effects of dysglycemia on neonatal development and known effects of maternal obesity on offspring metabolism [36–38] and brain development [3–7], only a few studies have begun to examine the effects of maternal obesity on glucose utilization in the offspring brain. Thus far, existing evidence suggest that offspring exposed to maternal obesogenic diet *in utero* decreased glucose uptake and utilization in the brain that can persist into adulthood [39,40]. Additionally, the vast majority of experiments have been done on mixed sex offspring, however, more recent studies have shown significant sex-differences in cerebral response to insults such as inflammation and hypoxia, with males being more susceptible [41]. The objective of our study was to understand the effects of maternal obesity on regulators of glucose transport and utilization in the fetal brain. We hypothesized that maternal obesity affects expression of key GLUTs involved in fetal brain glucose uptake, which results in cerebral energy deficits during fetal neurodevelopment.

## Methods

### Animals

The care and use of all mice were carried out according to an approved protocol by the Washington University Institutional Animal Care and Use Committee. C57BL/6J male and female mice were obtained from The Jackson Laboratory. At four-weeks-old, female mice were randomized to control chow (CON; 13% fat, 62% carbohydrates [3.2% sucrose], and 25% protein, PicoLab rodent diet 20) or an obesogenic chow (diet-induced-obesity, DIO: 14.9% protein, 59.4% fat, 25.7% carbohydrate [17.5% sucrose], Research Diets Inc diet D12451). Female mice had ad libitum access to food and water. Following 10 weeks on their respective diets, female mice were mated with male mice fed a standard chow. Timed mating was performed, pregnancy was confirmed by the presence of a vaginal plug on 0.5 days post coitum (dpc) and gestation was monitored. On 18.5 dpc, female mice were euthanized in a carbon dioxide chamber and fetal offspring were immediately extracted. Following fetal extraction, offspring were weighed, blood glucose levels were obtained (Akray Glucocard Vital Blood Glucose Monitoring System, Minneapolis, MN) and whole brains were isolated from the level of the olfactory bulb to the medulla. Whole brains were immediately flash frozen following extraction. One male and female litter pair were used per dam for subsequent analyses.

### Western Immunoblot

Whole brains were homogenized with lysis buffer containing 0.5% NP40 (ThermoFisher, MO, USA), 150 mM NaCl, 50 mM Tris-HCl (pH 7.5), 2% Glycerol, 1 mM EDTA supplemented with cOmplete Protease Inhibitor Cocktail (Roche, Indianapolis, IN) and Phosphatase Inhibitor Mini Tablets (ThermoFisher, MO, USA). Total protein was quantitated using Pierce’s BCA Protein Assay Reagent Kit (ThermoFisher, MO, USA) per the manufacturer protocol. 20 ug of protein from each sample were resolved in 4% to 12% Bolt Bis-Tris gels (ThermoFisher, MO, USA) and transferred to low-fluorescence PVDF membranes. Total protein concentration was estimated for GLUT protein quantitation prior to blocking the membranes with 5% nonfat dry milk by using Revert 520 Total Protein Stain Kit (LI-COR, NE, USA). Membranes were stained with antibodies against GLUT1 (1:5,000, Abcam Cat# ab115730) followed by Goat anti-Rabbit IgG (H+L) Highly Cross-Adsorbed Secondary Antibody, Alexa Fluor Plus 488 (1:10,000 ThermoFisher Scientific, Catalog # A32731), GLUT3 (1:2,500, Abcam Cat# ab191071) followed by Donkey anti-Rabbit IgG (H+L) Highly Cross-Adsorbed Secondary Antibody, Alexa Fluor Plus 800 (1:10,000 ThermoFisher Scientific, Catalog # A32808), and GLUT4 (1:2,000, Proteintech Cat# 66846-1-Ig) followed by Goat Anti-Mouse IgG Secondary Antibody, StarBright Blue 700 (1:2,500, BioRad Cat# 12004159). The signals were detected with fluorescence at 488nm, 700nm and 800nm wavelengths on a ChemiDoc MP (Bio-Rad Laboratories, CA, USA). Relative protein abundance was quantified using band signal to total protein within the same lane, normalized to the same first lane control for each blot using ImageLab (BioRad, Hercules, CA).

Phosphorylated AMPK and mTOR protein abundance was normalized to total AMPK and mTOR. Membranes were stained with antibodies against phosphorylated-AMPKα (Thr172) (40H9) (1:1,000 Cell Signaling Technology Cat# 2535) and antibodies against phosphorylated-mTOR (Ser2448) (1:1,000 Cell Signaling Technology, Cat# 2971S) followed by anti-rabbit IgG (H+L) HRP-conjugated secondary antibody (1:10,000 Jackson ImmunoResearch Labs Cat# 115-035-003) and imaged on the ChemiDoc MP (Bio-Rad Laboratories, CA, USA). After imaging, blots were then agitated at 55°C for 2 hours in a stripping solution of 0.5M Tris-HCl (pH 6.8), 10% SDS, washed with continuous deionized water, re-blocked in 5% nonfat dry milk and then re-probed with antibodies against AMPKα (1:1,000 Cell Signaling Technology, Cat# 2532S), and antibodies against mTOR (1:1,000 Cell Signaling Technology, Cat# 2972S) followed by anti-rabbit IgG (H+L) HRP-conjugated secondary antibody (1:10,000 Jackson ImmunoResearch Labs Cat# 115-035-003). The signal was detected with Supersignal West Femto (Thermo Fisher Scientific, Waltham, MA, USA) and the membranes were imaged on a ChemiDoc MP (Bio-Rad Laboratories, CA, USA). Relative protein abundance was quantified using phosphorylated band signal to total band signal ratio within the same lane, normalized to the same first lane control for each blot using ImageLab (BioRad, Hercules, CA).

### Sex Genotyping

Mouse tails were collected post-mortem for sex genotyping. DNA extraction was performed by heating samples to 95°C for 30 minutes in a 25 mM sodium hydroxide and 0.2 mM EDTA solution. Samples were then cooled to room temperature and neutralized with an equal amount of 40 mM Tris-HCl (pH 5.0) solution. Genomic DNA was amplified with the SX primer pair (SX_F, 5**′**-GATGATTTGAGTGGAAATGTGAGGTA-3**′** and SX_R, 5**′**-CTTATGTTTATAGGCATGCACCATGTA-3**′**) as previously described [42] and using Dream Taq Green PCR Master Mix (Thermo Fisher Scientific) as per the manufacturer protocol. The PCR parameters consisted of initial denaturation at 94°C for 3 min, 30 cycles with 94°C for 15 s, 60°C for 30 s, and 72°C for 15 s, followed by final elongation at 72°C for 3 min.

### Statistical Analysis

All statistics were performed using GraphPad Prism v10 software. Unless otherwise specified, significance was determined by two-way ANOVA with post hoc Holm-Sidak multiple comparison test, using the following cutoff values: \**P* < .05; \*\**P* < .01; \*\*\**P* < .001; \*\*\*\**P* < .0001.

## Results

### Maternal diet increases fetal circulating blood glucose levels

Our lab has previously shown that prolonged pre-pregnancy exposure of female C57BL/6J mice to an obesogenic diet induces metabolic changes pre-pregnancy and throughout gestation [43]. These include increased pre-pregnancy and gestational weight, insulin insensitivity, hypertriglyceridemia, and hyperlipidemia. Here, we confirmed that litter size was not affected by maternal diet (median litter size 8 pups [IQR 7-9] CON vs. 8 [IQR 3-10] DIO, p=0.97; Figure 1A). Offspring fetal weight at 18.5 dpc also did not differ by maternal diet with mean pup weight for CON offspring 1.010 ± 0.01 g compared to 1.013 ± 0.02 for DIO offspring (p=0.84; Figure 1B). However, fetal blood glucose levels were significantly higher in DIO offspring without modification by offspring sex (mean 45.6 ± 1.3 mg/dL CON vs. 53.4 ± 0.8 mg/dL DIO, p=0.02; Figure 1C). Finally, offspring whole brain weight (mean 63.6 ± 1.1 mg CON vs. 62.6 ± 0.7 DIO, p=0.2; Figure 2A) and fractional brain to body weight (mean 6.3 ± 0.07% CON vs. 6.2 ± 0.01% DIO, p=0.4; Figure 2B) did not differ by maternal diet.

**Figure 1:**
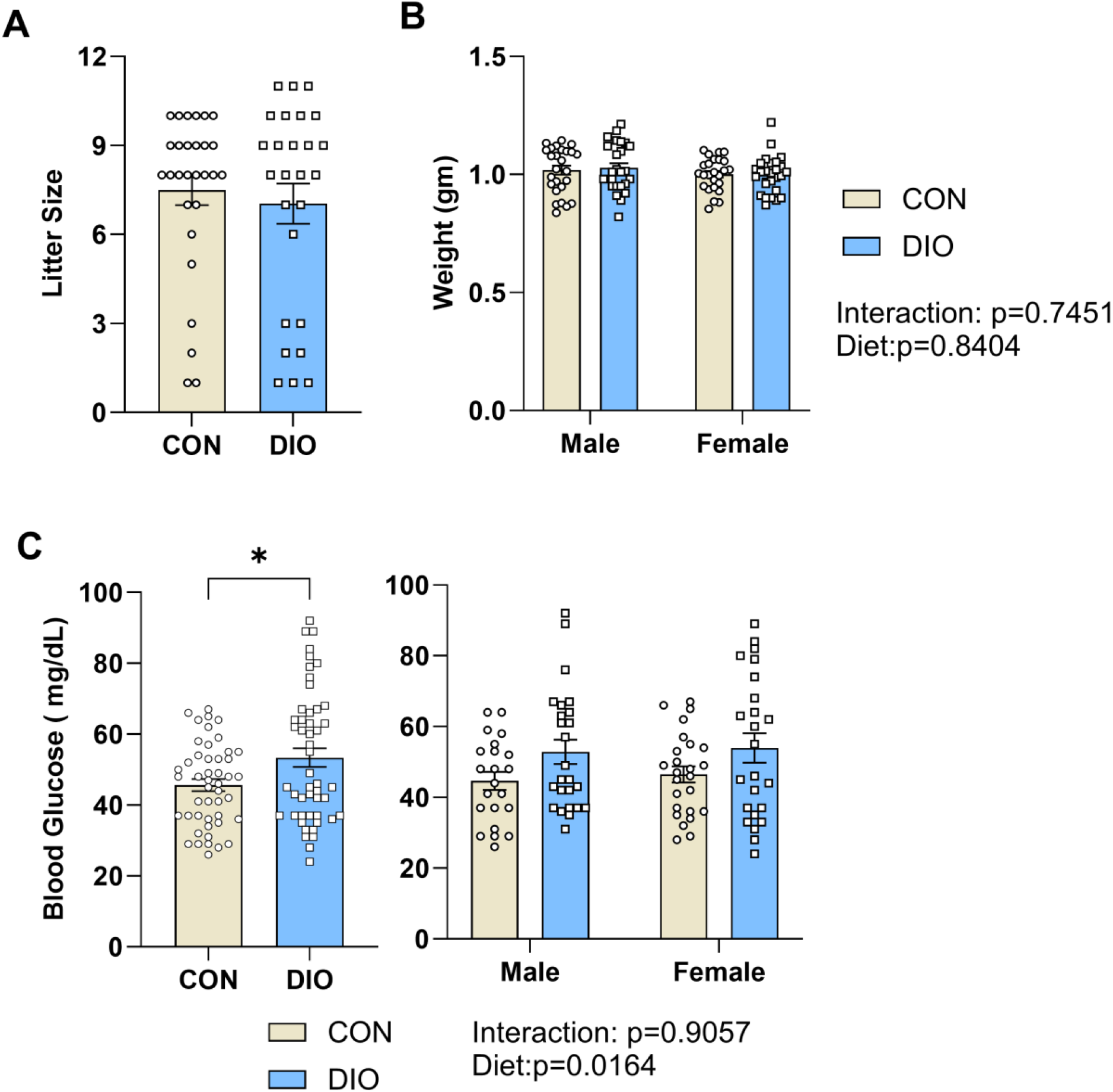
Fetal litter and offspring characteristics. There is no difference in (A) litter size or (B) offspring weight by maternal diet or fetal sex. (C) Fetal blood glucose was significantly higher among DIO offspring compared to CON and was not altered by fetal sex. Statistical significance for litter size were determined using the Mann-Whitney test. Statistical significance for offspring weight and blood glucose levels were determined using two-way ANOVA and adjusted for multiple comparisons using Sidak post hoc test (*p<0.05). CON, maternal control diet; DIO, maternal obesogenic diet.

**Figure 2:**
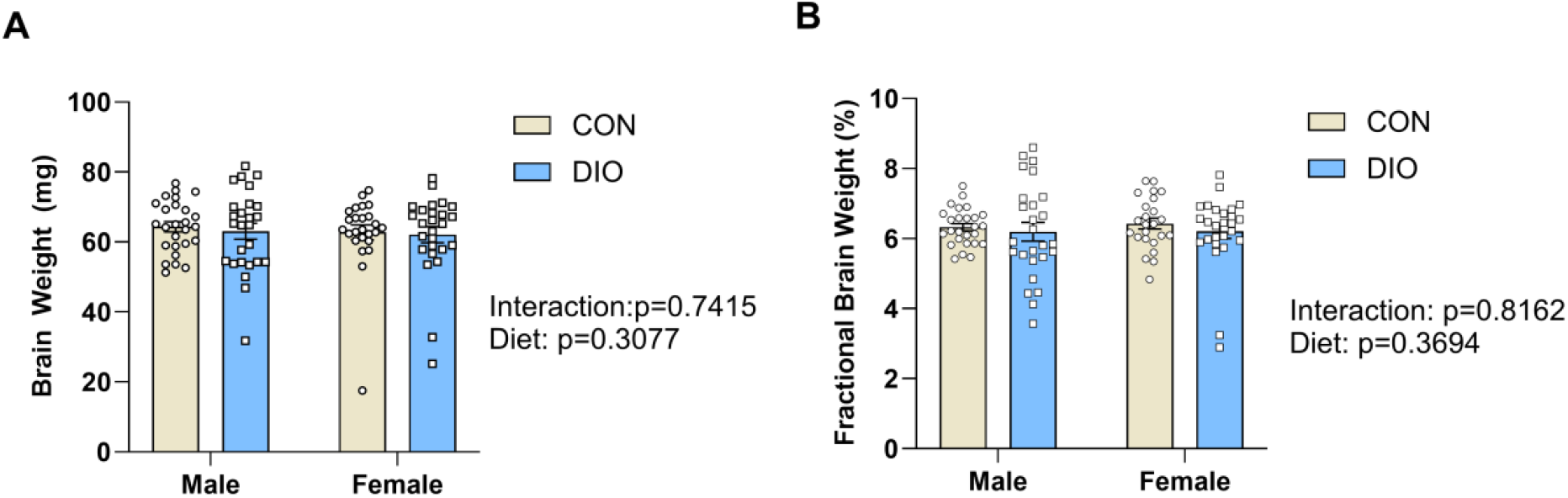
Fetal brain weight is not altered by maternal diet-induced obesity. (A) Fetal brain weight and (B) fractional brain to body weight does not differ CON and DIO groups. The individual data points on the graphs represent biological replicates. Error bars represent standard error of the mean. Statistical significance was determined using two-way ANOVA and adjusted for multiple comparisons using Sidak test. CON, maternal control diet; DIO, maternal obesogenic diet.

### Maternal diet increases fetal whole brain GLUT protein expression

GLUT 1 expression was higher in whole brain tissue isolated from female DIO compared to CON fetuses at 18.5 dpc (Figure 3A, C and D). However, the difference between sexes was not statistically significant in interaction modeling (p for interaction=0.13). Cerebral GLUT3 expression was higher in DIO offspring compared to CON (p=0.02) and this effect was not modified by offspring sex (p for interaction = 0.93; Figures 3B, C and D). Finally, abundance of the insulin-sensitive GLUT, GLUT4, was also higher in fetal whole brain from DIO compared to CON offspring (p=0.03), without modification by offspring sex (p for interaction=0.46; Figure 4).

**Figure 3:**
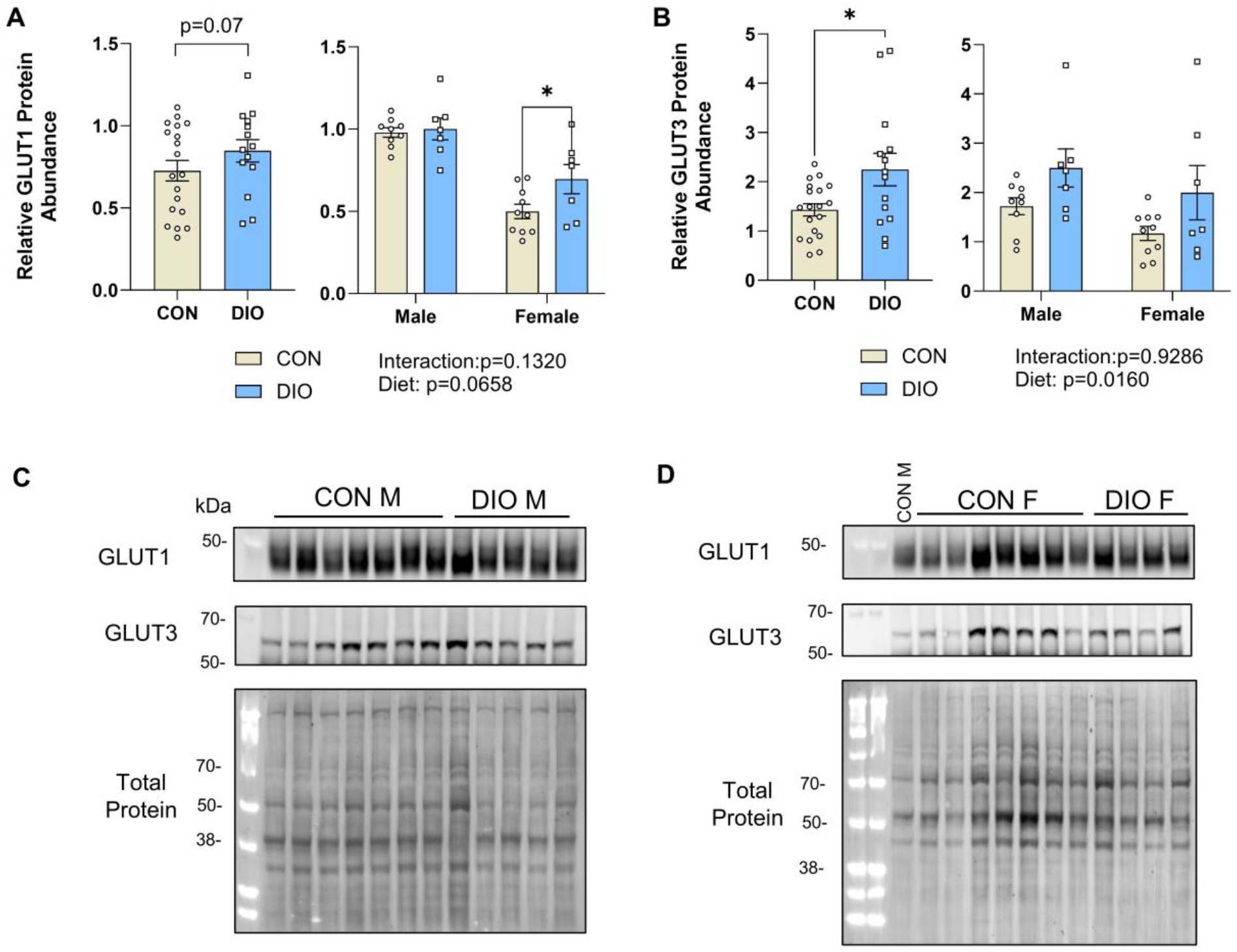
Maternal diet-induced obesity differentially alters whole brain GLUT expression in fetal offspring. (A) Quantification of fetal whole brain GLUT1 protein abundance normalized to total protein. (B) Quantification of fetal whole brain GLUT3 protein abundance normalized to total protein. (C and D) Representative western immunoblots are shown. The first lane in all blots (including the female blots) is a consistent control male sample and was used for normalization to allow for comparison between different blots. The individual data points on the graphs represent biological replicates. Error bars represent standard error of the mean. Statistical significance for relative protein abundance was determined using two-way ANOVA and adjusted for multiple comparisons using Sidak post hoc test (*p<0.05). CON, maternal control diet; DIO, maternal obesogenic diet.

**Figure 4:**
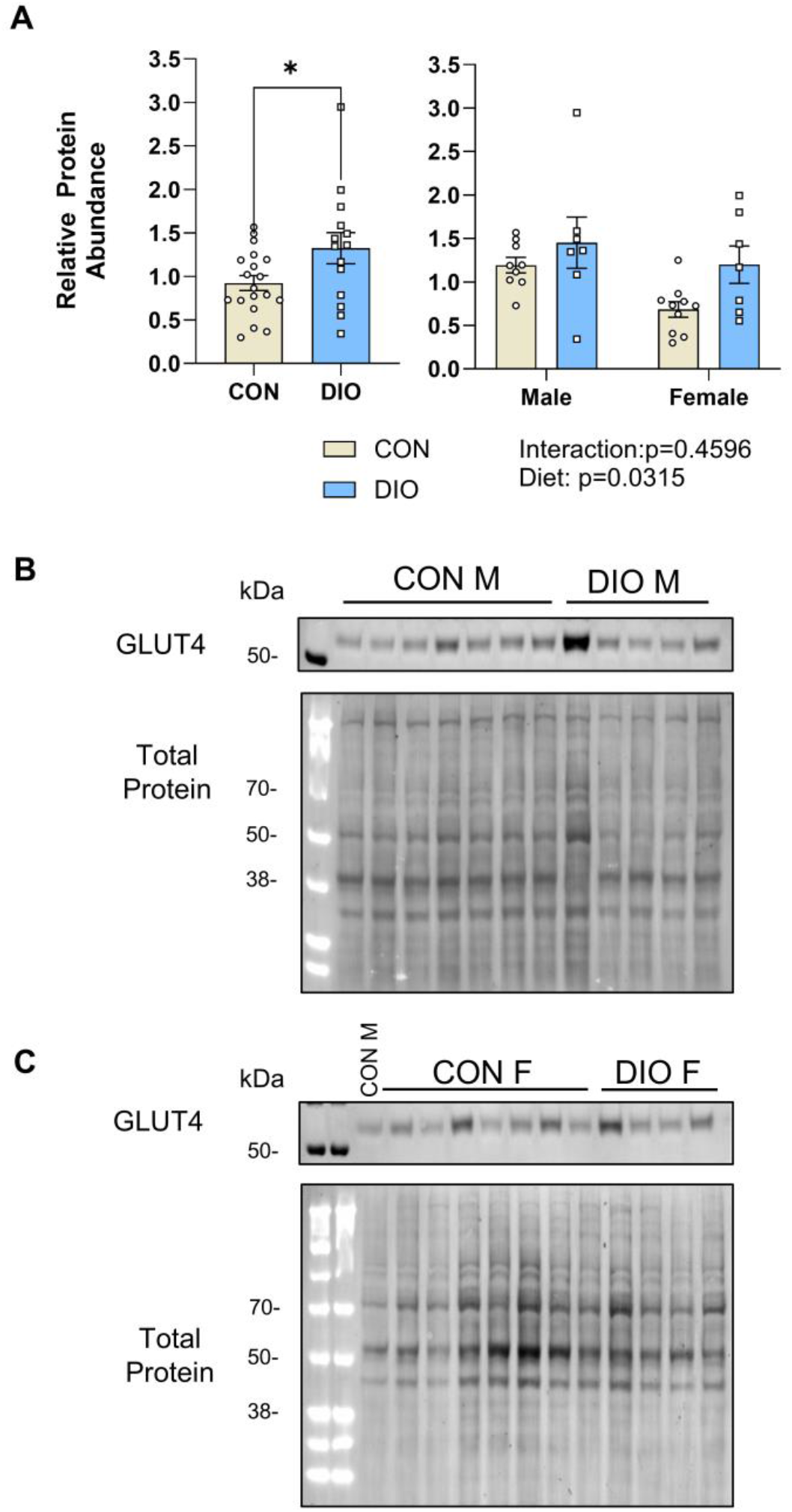
Maternal diet-induced obesity increases whole brain GLUT4 expression in fetal offspring. (A) Quantification of fetal whole brain GLUT4 protein abundance normalized to total protein and (B and C) representative western immunoblots. The first lane in all blots (including the female blots) is a consistent control male sample and was used for normalization to allow for comparison between different blots. The individual data points on the graphs represent biological replicates. Error bars represent standard error of the mean. Statistical significance for relative protein abundance was determined using two-way ANOVA and adjusted for multiple comparisons using Sidak post hoc test (*p<0.05). CON, maternal control diet; DIO, maternal obesogenic diet.

### Maternal diet does not contribute to fetal whole brain energy sensing dysregulation or growth

Given the observed differences in GLUT expression, we wanted to determine if the energy sensing pathways were differentially activated in the fetal brain of CON and DIO offspring. Adenosine monophosphate-activated protein kinase (AMPK) is a master energy sensor that primarily upregulates cellular catabolism [44]. Upon phosphorylation, AMPK is activated and acts on many substrates to increase both glucose transport and ATP production [45]. Mammalian target of rapamycin (mTOR) is another signaling pathway that can regulate cell metabolism and energy homeostasis and is negatively regulated by AMPK [46]. Our data show that neither phosphorylation of AMPK relative to total AMPK (Figure 5A, C and E) nor phosphorylation of mTOR (Figure 5B, D, F) differed by maternal diet in the fetal offspring brain.

**Figure 5:**
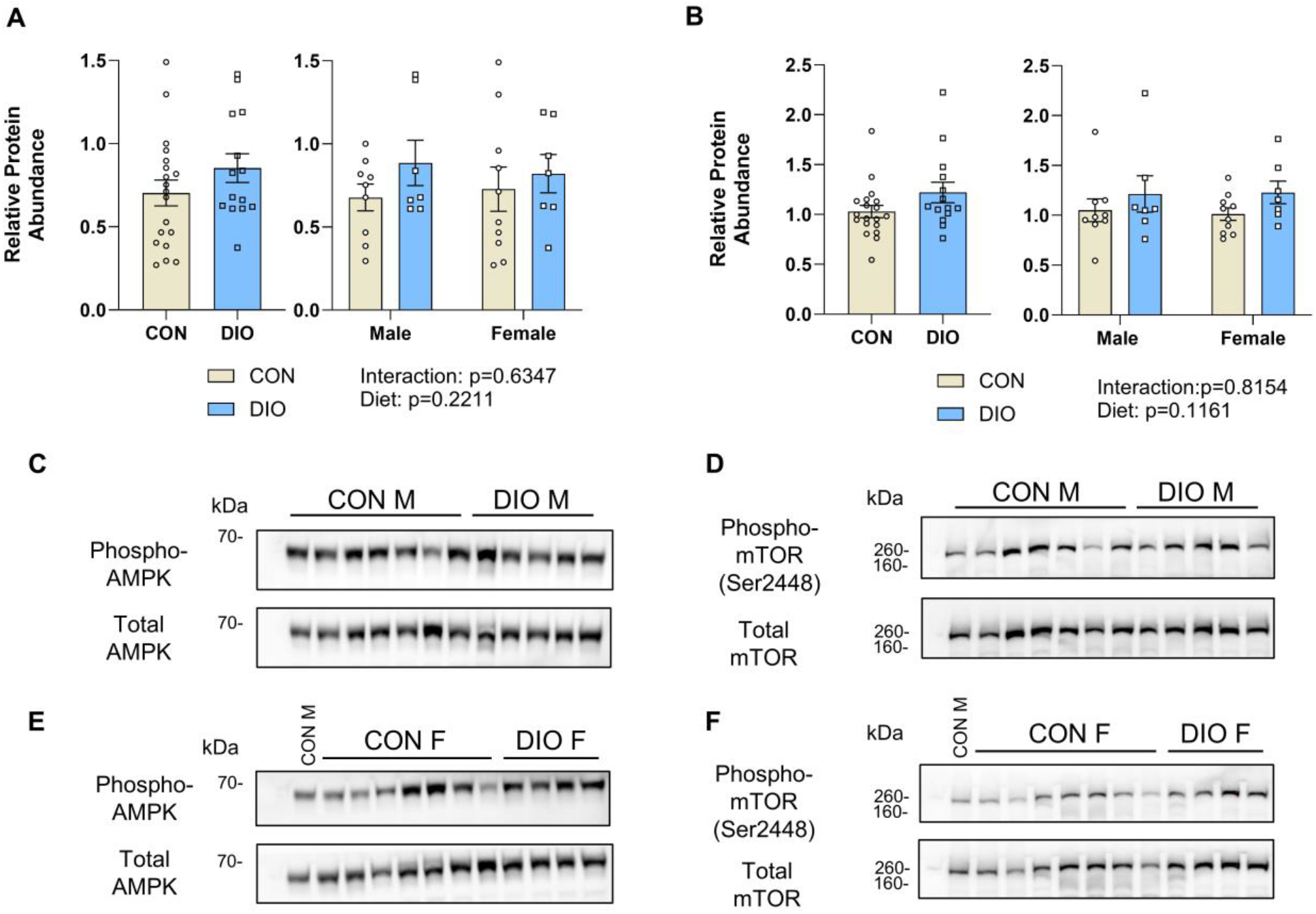
Fetal whole brain AMPK and mTOR phosphorylation are not altered by maternal diet. (A) Fetal whole brain phosphorylated AMPK is not different between CON and DIO offspring. (B) Fetal whole brain phosphorylated mTOR is not different between CON and DIO offspring. Representative western blots for (C and E) phosphorylated and total AMPK, and (D and F) phosphorylated and total mTOR, stratified by fetal sex. The first lane in all blots (including the female blots) is a consistent control male sample and was used for normalization to allow for comparison between different blots. The individual data points on the graphs represent biological replicates. Error bars represent standard error of the mean. Statistical significance for relative protein abundance was determined using two-way ANOVA and adjusted for multiple comparisons using Sidak test. CON, maternal control diet; DIO, maternal obesogenic diet.

## Discussion

Our current understanding of fetal brain metabolism, and particularly glucose uptake and metabolism, in the setting of maternal obesity is limited. This study suggests that maternal obesity affects GLUT protein expression in the fetal brain, particularly for neuronal GLUTs, but that the effect is modest. Additionally, there are no differences in activation of energy sensing pathways or fetal brain size, suggesting that exposure to maternal obesogenic diet in utero has minimal global effects on fetal cerebral glucose metabolism or perhaps that the observed GLUT upregulation is a balanced compensatory mechanism at this gestational age.

Alterations in fetal brain GLUT expression may predispose offspring to metabolic derangements during critical periods of neurodevelopment. Throughout pregnancy, glucose is freely transported across the placenta, and fetal supply is directly proportional to maternal blood glucose concentrations [47]. This means that in the setting of maternal hyperglycemia, such as with metabolic syndrome, the fetus is also exposed to a hyperglycemic environment. This exposure is believed to contribute to the pathogenesis of fetal growth abnormalities observed in patients with diabetes mellitus [48]. The effects of in utero hyperglycemia on fetal neurodevelopment are less well characterized; however, evidence can be gleaned from adult and neonatal studies. Abnormalities in glucose supply and transport in the adult brain can influence the outcome of neurodegenerative disorders such as Alzheimer’s disease and Parkinson’s disease, stroke, and traumatic brain injury [45]. Glucose dysregulation in the neonatal period can also impair cerebral development. For example, episodes of neonatal hypoglycemia are associated with lower executive and visual motor function in preschool-aged children [35]. In premature infants, prolonged hyperglycemia is associated with impaired neurodevelopment at 12 months of age [49]. Moreover, in cases of moderate to severe hypoxic ischemic encephalopathy, both hyperglycemia and hypoglycemia have been associated with abnormal neurodevelopment or death at 18 months of age [50]. Several animal studies have also shown that in utero exposure to maternal obesity affects glucose utilization in the offspring brain. The strongest evidence for long-term effects comes from Sanguinetti et al., which showed that offspring of minipigs exposed to a high-fat diet in utero exhibited decreased baseline glucose uptake and glycogen deposition in the brain following birth and persisting for at least 12 months [40]. Another study showed that glucose utilization is decreased in the hypothalamus of offspring of rats with diet-induced obesity, but did not examine other cerebral tissues, developmental timepoints or mechanisms [39]. While our study did not specifically examine glucose uptake in the fetal brain, our findings suggest an increase in glucose transporter expression, which may be compensatory and prevent energetic deficiency.

Our data show that fetal whole brain GLUT3 and GLUT4 expression are increased in the setting of maternal obesity without modification by sex. In rodents, neuronal GLUT3 and GLUT4 expression is upregulated in the early postnatal period, coincident with increased synaptogenesis and maturation [51–53]. The increase in fetal brain GLUT3 and GLUT4 expression in animals exposed to DIO therefore may indicate increased glucose need during this developmental period. Previous studies suggest that GLUT expression and uptake are regulated by circulating glucose levels [54,55]. GLUT3 is a high affinity for glucose and is more efficient in glucose transport than GLUT1 [56]. In the neuron, GLUT3 is highly expressed at pre and post-synaptic nerve endings suggestive of its role in energy demanding processes such as nerve conduction [14]. Therefore, increased GLUT3 expression in the brain may signify increased glucose utilization or impaired glucose delivery. Adult rodent high fat diet (HFD) and diabetic models have resulted in varying effects on GLUT3 expression, which may reflect specific tissue needs. Despite decreased global cerebral glucose uptake, whole brain GLUT3 expression did not differ in an adult diabetic mouse model [57]. In another study, hypothalamic GLUT3 expression was decreased and hippocampal GLUT3 expression showed no change after exposure to a HFD [58]. These differences may be explained by varying pathologic responses to experimental design. Diabetic phenotypes are often induced with streptozotocin, thus creating an insulin deficient animal model of diabetes. This leads to a phenotype characterized by hyperglycemia and ketoacidosis, both of which can contribute to mechanistic changes in cerebral glucose transport [59]. Depending on the duration of a HFD exposure, insulin resistance can occur and diabetic phenotypes may contribute substantially to pathophysiologic changes. In the current study, there was some evidence of maternal insulin resistance from gestational GTTs, though, we do not suspect the mild elevation in blood glucose levels observed was sufficient to induce insulin resistance in the fetus.

Similar to GLUT3, GLUT4 expression is upregulated in brain regions with high energetic demands but can also be regulated by insulin signaling [34]. GLUT4 is primarily located in neurons and often colocalizes with GLUT3 and the insulin receptor [14]. In the axonal membranes, GLUT4 has also been shown to be regulated by AMPK, an energy sensing pathway activated in decreased energy states [33]. Sustained neuronal activation increases GLUT4 expression into the axonal plasma membrane supporting its regulation in metabolically active cell types. In this experiment, the increased fetal brain GLUT4 in DIO offspring likely occurred from a combination of insulin signaling and increased metabolic demand.

Fetal cerebral GLUT1 has an important role in glucose supply during early brain development [15,60]. GLUT1 expression in the fetal period is primarily at the endothelial cells of the blood brain barrier and dendritic processes of surrounding astrocytes, albeit at levels much lower than that of the adult [53,61]. Prior studies have shown increased GLUT1 expression in hippocampal and hypothalamic glial cells in animals exposed to overnutrition in the neonatal period [62]. In this study, we did not see an increase in whole brain GLUT1 expression in the fetal offspring. This could be due to blunting of region-specific effects in the setting of whole brain analysis or could signify a decreased role for this receptor in cerebral glucose homeostasis during fetal compared to postnatal development [63]. Additionally, although interaction by sex did not reach statistical significance, we did detect a significant increase in GLUT1 expression among female offspring exposed to DIO in our stratified analyses, suggesting that overnutrition insults can have sex-specific effects. Earlier studies excluded female offspring in experimental design and therefore our understanding of sex differences in dietary effects on neurodevelopment are limited [8]. However, sex differences in response to cerebral insults such as traumatic brain injury, cerebral hypoxic-ischemic injury, and cerebral palsy have previously been shown, but usually show higher susceptibility among males [64–66]. Additionally, studies examining effects of high fat diets on neurological outcomes such as neurogenesis or susceptibility to dementia have suggested a greater effect in females [67,68]. Alternatively, other rodent studies have shown decreased glucose uptake in male brains exposed to overnutrition. Therefore, it is also possible that the increased GLUT1 expression seen in the female offspring has a protective effect and is a mechanistic explanation for the higher susceptibility to various cerebral insults seen among male offspring.

It is important to note that two isoforms are commonly seen for the GLUT1 receptor, a larger endovascular transporter and smaller glial cell transporter [14,56]. Vascular GLUT1 is larger because it has varying degrees of post-translational modification including glycosylation [51,69]. In this study, we did not distinguish distinct bands to account for vascular versus glial cell GLUT1 protein expression, but saw a continuous diffuse band visualized at 45-50 kDa on western immunoblot, which likely representative of a combination of glial and endovascular GLUT1 expression. As a result, a more profound difference in cell-specific GLUT1 expression, may not have been appreciated in our analyses.

We evaluated the AMPK and mTOR pathways to understand whether energy availability was affected in the DIO fetal offspring brains. AMPK is a heterotrimeric complex activated by the presence of adenosine monophosphate (AMP) [70]. AMPK phosphorylation and activation occurs in response to either a decreased energy environment or by cellular stress from reactive oxygen species [71,72]. Downstream effects of this pathway are to maintain homeostasis and restore energy rich ATP through regulation of catabolic and anabolic processes [70]. Decreased hypothalamic AMPK phosphorylation has been reported in postnatal day 1 mouse offspring exposed to a maternal high fat diet likely from an increased energy level state [73]. Additionally, neuronal oxidative stress may contribute to decreased neuronal glucose uptake despite normal to increased GLUT expression [74]. Mechanistic target of rapamycin (mTOR) is a highly conserved protein kinase that controls nutrient status and cell growth [75]. Upregulation of mTOR is associated with increased nutrient delivery and in maternal obesity has been associated with fetal overgrowth [76]. We did not see an effect of maternal diet on fetal whole brain AMPK or mTOR activation. This suggests both an appropriate energy balance as well as minimal to no cellular oxidative stress. We postulate that this may reflect a balanced compensatory effect of observed cerebral GLUT expression in the DIO offspring. This possibility is additionally supported by our data showing no differences in brain weight.

While our lab has previously shown that an obesogenic diet altered the metabolic profile of female mice prior to pregnancy and duration gestation [43], we did not see a difference in fetal weight by maternal diet at 18.5 dpc. Prior rodent studies show conflicting data with some groups showing an increase in male offspring birthweight [65], while others report a decrease in fetal weight at 14.5 dpc [66]. The high variability in outcomes between studies likely reflects the significant variation in experimental design when assessing offspring weight by maternal diet. Differences in mouse strain, diet macronutrient composition, diet exposure duration and offspring age at assessment can all affect outcomes [77].

Blood glucoses were significantly increased in fetal DIO offspring. This may be in part from a combination of maternal insulin resistance and placental regulation of glucose transport. Our lab has previously reported that gestational DIO mice showed some evidence of insulin insensitivity by oral gavage glucose tolerance tests [43]. Although we did not directly evaluate this, glucose transport in the placenta occurs primarily through GLUT1 [12] and it has been shown that placental GLUT1 expression in the basal membrane is increased with maternal obesity [78]. Therefore, increased circulating glucose during gestation and increased glucose transport capacity through placental GLUT1 expression may account for elevated fetal blood glucose levels.

There are several limitations to this study that must addressed when interpreting the data presented. First, the obesogenic diet we used contains approximately 60% of calories from fat, and this is primarily saturated fat. This contrasts with the standard Western diet reporting an average fat and carbohydrate consumption between 30-35% and 45-50%, respectively [79]. Additionally, whole brain samples were analyzed for quantification of GLUT protein expression, as well as AMPK and mTOR activation. As a result, there may be significant differences within specific brain regions that were not appreciated. Differences in energy sensing may have been present in regions with high energy demands, such as the hypothalamus of the developing fetal brain, and missed on whole brain analysis. We also only analyzed the fetuses at one time point of late gestation and did not assess temporal differences. Finally, this study did not evaluate expression of other GLUTs, including GLUT2, which has been reported in the brain [80]. Functionally, it is involved in the regulation of glucose uptake and, within the brain, is localized primarily to the hypothalamus [14]. Future work should focus on characterizing the spatiotemporal expression of GLUTs and in offspring brains throughout in utero and postnatal development. Additionally, functional characterization of cerebral metabolism, including glucose uptake, should be assessed in order to correlate with our findings of differential GLUT expression in the setting of maternal obesogenic diet exposure.

## Conclusions

The maternal obesity epidemic continues to impact maternal-fetal health and may be a significant contributor to fetal neurodevelopmental programing. Using a validated maternal obesity murine model resulted in both maternal and fetal phenotypic changes. Specifically, diet induced obesity altered the maternal metabolic profile and fetal offspring showed hyperglycemia.

Our findings suggest that maternal obesogenic diet alters glucose transporter expression in the fetal brain, with the greatest effect on neuronal GLUTs 3 and 4. These differences may affect glucose uptake and metabolism during critical periods of perinatal development. Further studies in fetal cerebral glucose metabolism in the setting of maternal obesity are necessary to elucidate whether the observed changes in GLUT expression contribute to fetal and neonatal neurodevelopment.

## Abbreviations

SLC2: solute carrier 2
GLUT: glucose transporter
CON: Control
DIO: obesogenic
dpc: days post coitum
HFD: High Fat Diet
AMPK: adenosine monophosphate activated protein kinase
AMP: adenosine monophosphate
mTOR: mechanistic target of rapamycin
GTT: glucose tolerance test

## Acknowledgements

This work is in part supported by the National Institute of Child Health and Human Development (R21 HD110610 to A.I.F.).

